# Non-linear phenotypic variation uncovers the emergence of heterosis in *Arabidopsis thaliana*

**DOI:** 10.1101/404616

**Authors:** François Vasseur, Louise Fouqueau, Dominique de Vienne, Thibault Nidelet, Cyrille Violle, Detlef Weigel

## Abstract

Heterosis describes the phenotypic superiority of hybrids over their parents in traits related to fitness. Understanding and predicting non-additive inheritance such as heterosis is crucial for evolutionary biology, as well as for plant and animal breeding. However, the physiological bases of heterosis remain debated. Moreover, empirical data in various species have shown that diverse genetic and molecular mechanisms are likely to explain heterosis, making it difficult to predict its emergence and amplitude from parental genotypes alone. In this study, we evaluated a model of physiological dominance proposed by Sewall Wright to explain the non-additive inheritance of metabolic fluxes at the cellular level. We used 450 hybrids derived from crosses among natural inbred accessions of *Arabidopsis thaliana* to test Wright’s model for two fitness-related traits at the whole-plant level: growth rate and fruit number. We found that allometric relationships between traits constrain phenotypic variation in hybrids and inbreds to a similar extent. These allometric relationships behave predictably, in a non-linear manner, explaining up to 75% of heterosis amplitude, while genetic distance among parents at best explains 7%. Thus, our findings are consistent with Wright’s model of physiological dominance on plant performance, and suggest that the emergence of heterosis is an intrinsic property of non-linear relationships between traits. Furthermore, our study highlights the potential of a geometric approach of phenotypic relationships for predicting heterosis of two major components of crop productivity and yield.

## Introduction

If the inheritance of phenotypic traits was always additive, progenies will always exhibit intermediate trait values compared to their respective parents. Pure genetic additivity is, however, the exception rather than the rule in intraspecific crosses. Non-additive inheritance has been exploited for decades in agronomy [1,2], although the underlying mechanisms remain a major question for evolutionary genetics and crop science [2,3]. In plants, hybrid vigour or heterosis has been frequently observed [4–9], and its molecular bases have been investigated in numerous studies [9–13]. Unfortunately, empirical observations often led to contrasting conclusions, and none of the proposed genetic mechanisms can fully explain the emergence of heterosis of different traits across all study systems [9,14–19]. Thus, we are still lacking a unifying theoretical corpus that enables us to explain and predict heterosis of fitness-related traits, including biomass, growth rate and reproductive success.

Several genetic hypotheses have been proposed to explain heterosis [3,10,13,20]. According to the ‘dominance hypothesis’, each parent contributes favourable, dominant alleles (generally at many loci) that together complement the deleterious effects of recessive alleles originating from the other parent. The ‘overdominance hypothesis’ postulates the existence of loci where the heterozygous state (*Aa*) is superior to both homozygotes (*AA* or *aa*). With overdominance, the emergence of a hybrid superior to its best parent could depend on a single locus. Pseudo-overdominance corresponds to cases where dominant, favourable alleles are in linkage with recessive, unfavourable alleles, so that the heterozygous combinations appear to behave as overdominant loci. The picture is further complicated by the contributions of epistasis [15,21] and epigenetics [12,22] to heterosis in plants.

To tease apart the different hypotheses, quantitative trait locus (QTL) mapping has been carried out in many species [5,9,15,17,21,23]. In general, individual studies differ strongly in their conclusions regarding the underlying genetic mechanisms, apparently depending on the investigated traits, the genetic material used, the mating system, and the experimental constraints related to the number of crosses necessary for robust statistical inference (*e.g.*, diallel mating design) [13,19,20]. In addition, the QTL approach performs poorly to detect small-effect loci [24]. Yet, it has been shown that fitness-related traits such as growth rate, size and fruit production are often controlled by a large number of genes, which individually may have very weak effects [25–27].

Because of the polygenic nature of fitness-related traits, heterosis is expected to be associated with molecular dominance, and should positively correlate with genetic distance between parents, at least up to a certain extent [28–30]. Some findings are consistent with this hypothesis [31–33], suggesting that parental genetic distance could be used to quantitatively predict heterosis in plants. Unfortunately, experimental studies generally employ for practical reasons only a relative small number of crosses and parental lines [7,34], which makes it difficult to generalize the findings of individual studies. In recent work, Seymour and colleagues [13] investigated heterosis in a half-diallel between 30 genome-sequenced accessions of *Arabidopsis thaliana* collected from diverse Eurasian populations [35]. As expected under the dominance model [28–30], they found a positive correlation between parental genetic distance and heterosis. However, genetic distance between parents only accounted for less than 3% of heterosis among *A. thaliana* hybrids, making predictions based on genetic distances alone strongly uncertain.

Strikingly, many continue to consider the physiological bases of heterosis a mystery [36,37]. An alternative to genetics-first studies in order to understand and predict heterosis is to consider the physiological constraints that determine phenotypic variation across organisms. As early as 1934, Sewall Wright proposed a model of physiological dominance to explain the maintenance of recessive alleles in natural populations [38]. Wright began with the universal relationship that connects the concentration of enzymes to the metabolic flux that results from their activity (Fig. 1). Since the relationship between these two traits is concave with a horizontal asymptote (*e.g.*, Michaelis-Menten kinetics), dominance of metabolic flux is expected even if enzyme concentrations are additive and hybrids intermediate between their parents (Fig. 1). Modern knowledge of metabolic networks was incorporated into this model by Kacser and Burns in 1981 [39], but Wright’s model of dominance has otherwise received little attention in the heterosis literature. Recently, Fiévet and colleagues [40,41] evaluated Wright’s model with a focus on the relationship between metabolic flux and enzyme activity in the chain of glycolysis in yeast. They modelled the importance of the curvature of the relationship between enzyme concentration and metabolic flux in the emergence of heterosis on glycolysis output [40]. This approach has considerable promise for predicting the phenotype of hybrids by considering the non-linear relationships that often link phenotypic traits with each other. However, these theoretical expectations remain to be tested on more complex traits, as well as on multicellular organisms such as plants, with potentially major perspectives for cultivated species.

**Figure 1.**
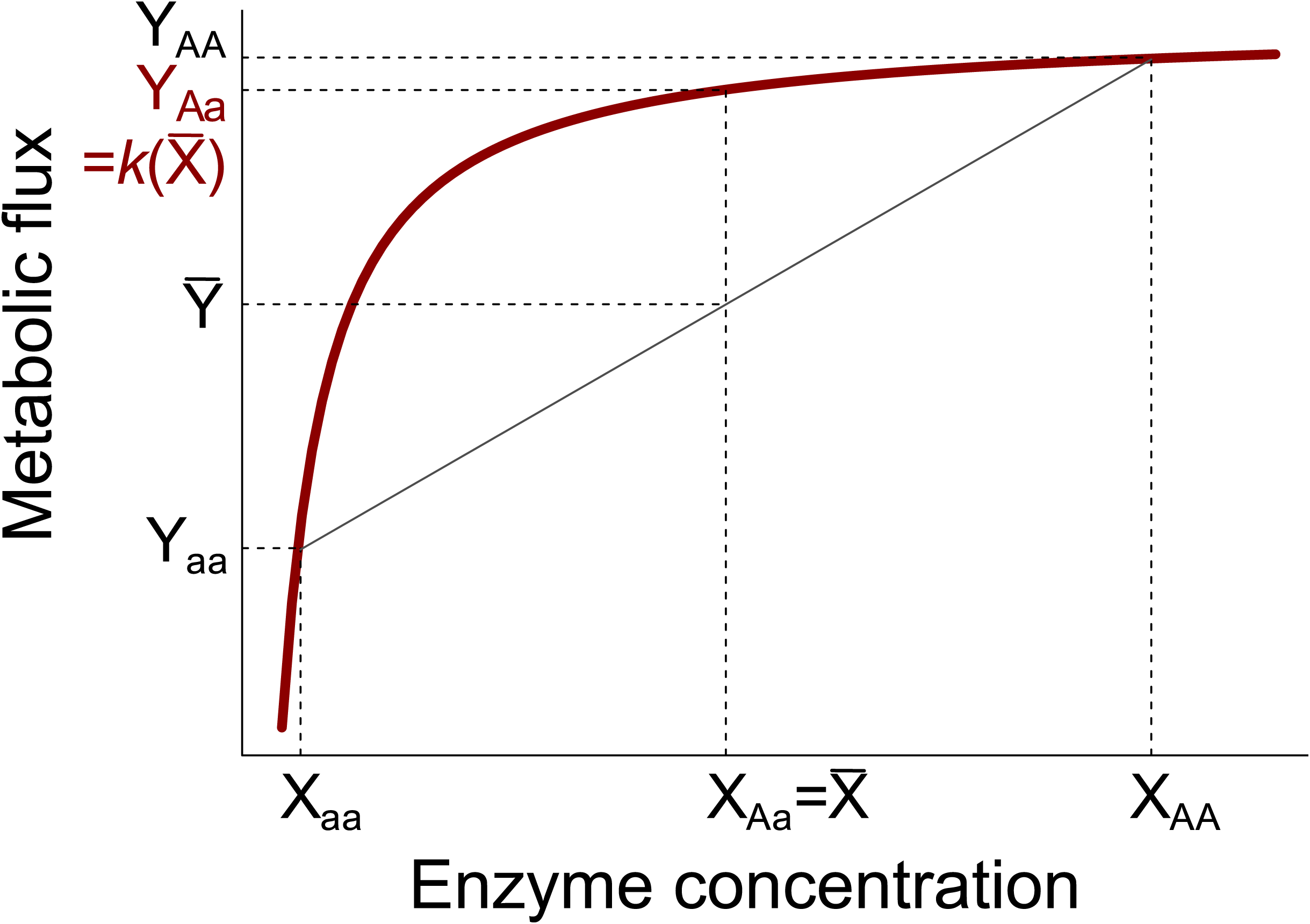
Wright’s model of physiological dominance. The metabolic flux along the y axis exhibits heterosis as a result of non-linear relationship with the concentration of enzymes along the x axis. This is expected for hybrids (genotype *Aa*) even if the concentration of enzymes is purely additive, as shown here (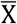 is mean enzyme concentration value between parent 1 (*aa*) and parent 2 (*AA*)). Red line represents Michaelis-Menten kinetics of enzyme activity (*k*(X)).

Many phenotypic relationships exhibit non-linear geometries, notably between fitness-related traits [42,43]. These relationships are analogous to those between metabolic and enzymatic activities described by Wright [38], Kacser and Burns [39], and Fiévet and colleagues [40,41]. In particular, comparative ecology studies have demonstrated the existence of non-linear allometric relationships between the biomass of an organism and its morphology, physiology and metabolism [42,44–46]. Allometric relationships often take the form of a function Y=αM^β^, where Y is a morphological or physiological trait (*e.g.*, respiration rate, growth rate, leaf area, reproductive biomass) that can be predicted by *M*, the biomass of the organism, a constant α and the allometric exponent β [44,47]. The allometric exponent of many traits exhibits a value less than one, causing a concavity of the relationship (called ‘strict allometry’ [42]). Allometric relationships are usually linearized by logarithmic transformation, especially to compare exponent values (slope of the log-log relationship) between organisms [48]. For instance, the metabolic scaling theory predicts that the allometric exponent is a multiple of a quarter (e.g., ¼, ¾) for many biological processes such as respiration rate, hydraulic resistance, carbon fluxes, biomass allocation and growth rate [44,46,49–51]. Recent studies with plants [43,52] suggest that the exponent must be corrected for the organism’s biomass because it is linked to the adaptive diversification of plant metabolism [27].

In this study, we tested, inspired by Wright’s model of physiological dominance for enzyme concentration and metabolic flux [38], whether non-linear allometric relationships can explain the emergence of heterosis in macroscopic traits related to performance, such as growth rate and fruit production. We used the plant *A. thaliana*, which is not a crop, but has been used for many genetic studies of heterosis [7,13,53–56]. This species is characterized by a high rate of inbreeding, which leads to a high rate of homozygosity in natural populations [57,58]. The complete sequencing of 1,135 natural accessions [59] has provided abundant information on the extent of genetic variation between populations, the genotype-phenotype map of the species, as well as the causes of phenotypic variation in intraspecific hybrids [60–64]. Furthermore, allometric relationships that link growth rate and fruit production to plant biomass have been evaluated in this species [27,52,65]. Here, we specifically addressed the following two questions: (i) Are there non-linear allometric relationships that similarly constrain trait variation in natural accessions and hybrids? And (ii) do these non-linear relationships explain the emergence and extent of heterosis in growth rate and fruit production? To this end, we compared several traits between 451 inbred accessions representing a wide range of native environments and 450 crosses derived from crosses among 415 of these accessions. Our results are consistent with many aspects of Wright’s model, suggesting that heterosis emerges intrinsically from the non-linear relationships between traits.

## Results

### Trait variation and heterosis in a wide range of *A. thaliana* genotypes

We generated 450 hybrids by manual crosses between 415 inbred accessions (Fig. 2A). We then measured four traits, vegetative dry mass, plant lifespan, growth rate and total fruit number, in the 450 hybrids, the 415 inbred parents, plus a further 35 inbred accessions. The parental combinations for the hybrids had been chosen to represent a wide range of genetic, geographic and phenotypic distances (Fig. 2B,C).

**Figure 2.**
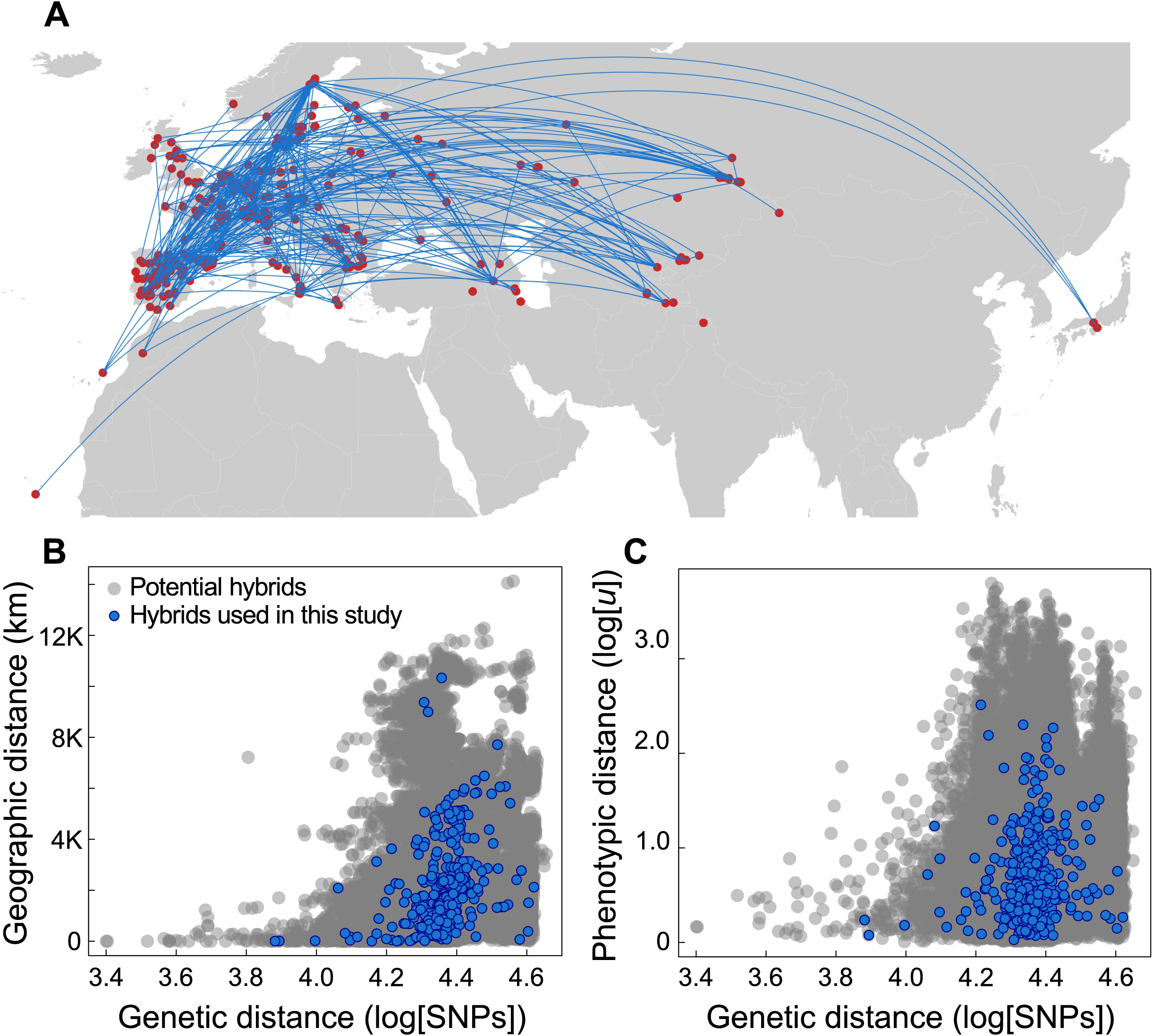
Diversity of genetic crosses used. (**A**) Origin of 451 natural inbred accessions (red dots), and the 450 hybrids between them (blue connecting lines). (**B**) Relationship between pairwise genetic distances and pairwise geographic distances. (**C**) Relationship between pairwise genetic distances and pairwise phenotypic distances (Euclidean distances). Distance-distance relationships are represented between all possible pairs of the 451 accessions (101,475 combinations, grey dots) and the crosses used for the 451 phenotyped hybrids (blue dots).

Traits were strongly variable between genotypes: vegetative dry mass *M* varied between 1.3 and 2,218 mg, plant lifespan between 24 and 183 days, growth rate between 0.04 and 40.4 mg d^−1^, and fruit number between 5 and 400 (Table S1). Across the entire dataset of hybrids and inbreds, a high proportion of the phenotypic variance could be explained by genotypic differences (broad-sense heritability): from 80% for growth rate to 94% for vegetative dry mass (Table S2). We found similar ranges of variation for both inbreds and hybrids (Fig. 3). For instance, standard deviation of plant lifespan was 19.8 days across inbreds, and 20.1 days across hybrids (Table S2). Hybrids were on average not significantly different from inbreds (*P* > 0.01 for all traits, Table S2).

**Figure 3.**
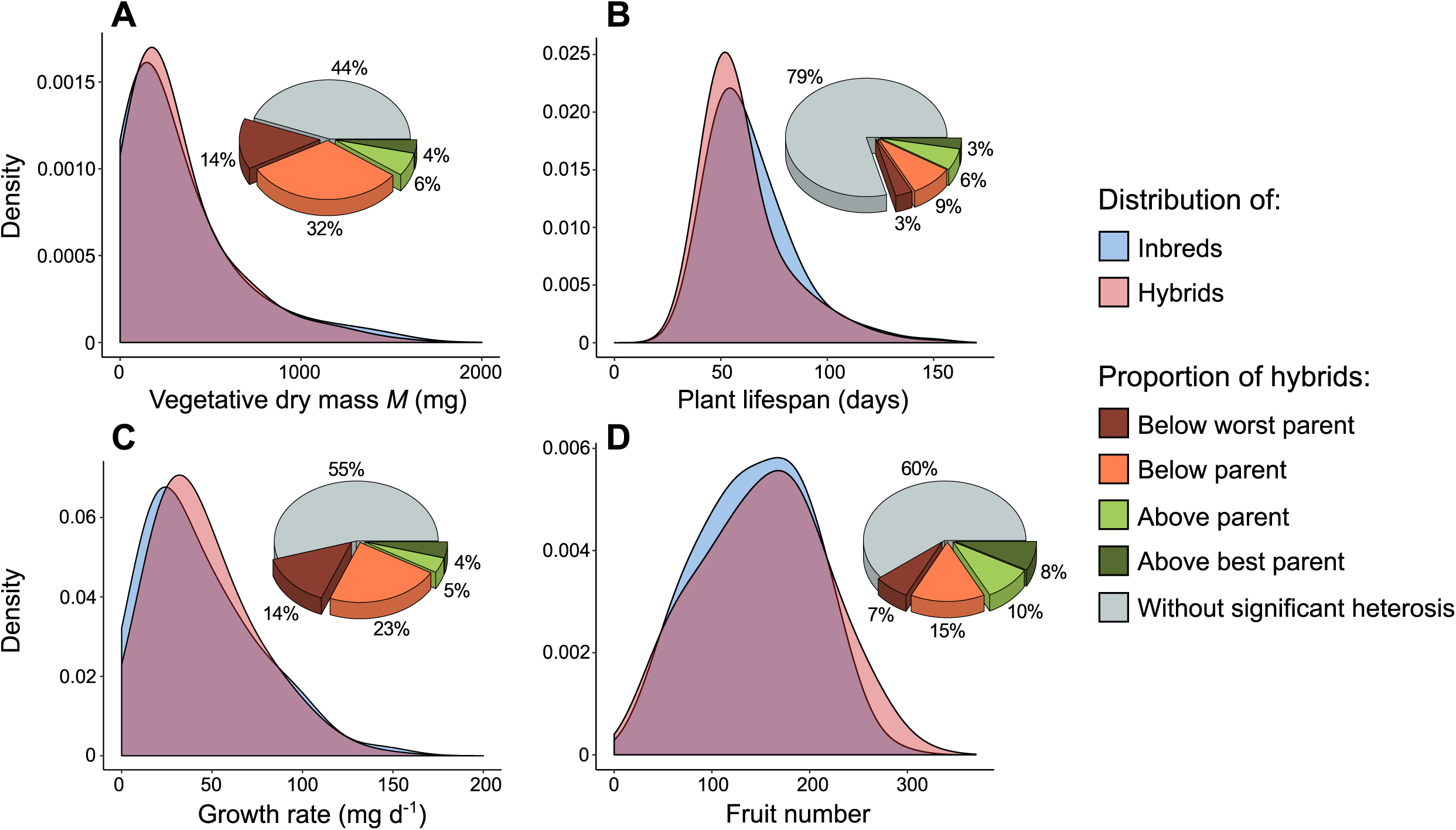
Distributions of traits and proportion of heterosis. (**A**) Vegetative dry mass *M* in inbreds (red) and hybrids (blue). Pie chart indicates proportion of crosses without and with significant heterosis. (**B**) Plant lifespan. (**C**) Growth rate. (**D**) Fruit number.

Despite the similar distribution of trait values for inbreds and hybrids, we found significant heterosis for all traits. This was measured by the discrete categorization of hybrid phenotypic class based on the comparison of hybrid trait distribution with both mid-parent values and highest (‘best’) or lowest (‘worst’) parent values. There was both positive and negative heterosis (Fig. 3). However, the extent and direction of heterosis was variable between traits. For instance, 21% of hybrids had confidence intervals (CIs) for lifespan that significantly differed from both or mean parental value, and thus, were considered as significantly heterotic. By contrast, 56% of hybrids were heterotic for vegetative dry mass. In general, there was more negative than positive heterosis, specifically for vegetative dry mass (46% of hybrids). Thus, hybrids represent a similar phenotypic space as natural inbred accessions, and the emergence of heterosis in our data did not simply reflect a few exceptional hybrids that outperformed all inbreds.

### Similar non-linear allometric relationships in accessions and hybrids

In this study, we focused on two traits with important evolutionary and agronomic outcomes: growth rate and fruit number. As predicted by ecological theory, these two traits exhibited non-linear allometric relationships with vegetative dry mass (Fig. 4). Consistent with recent studies of plant allometry [27,43,52], growth rate could be modelled by a power-law function with mass-corrected exponent (*g*(*M*), Table 1), with a concave curvature (Fig. 4A). By contrast, fruit number exhibited a right-skewed bell-shaped relationship (Fig. 4B). We modelled this allometric relationship with an inverse quadratic function (*f*(*M*), Table 1). Importantly, inbreds and hybrids exhibited similar non-linear relationships, characterized by coefficients that were not significantly different from each over (with overlapping 95% CIs; Table 1).

**Table 1.**
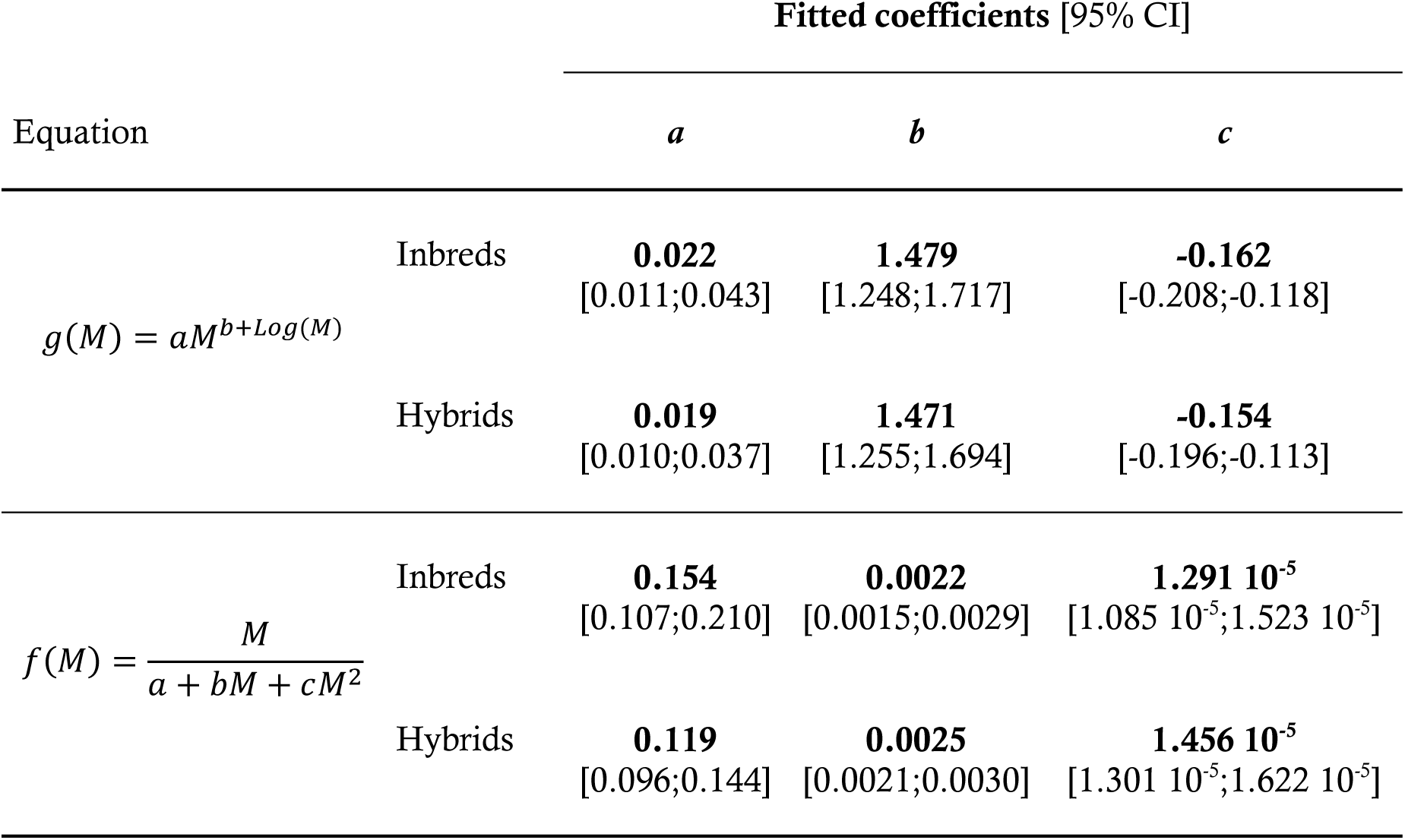
Coefficients of fitted allometric relationships.

**Figure 4.**
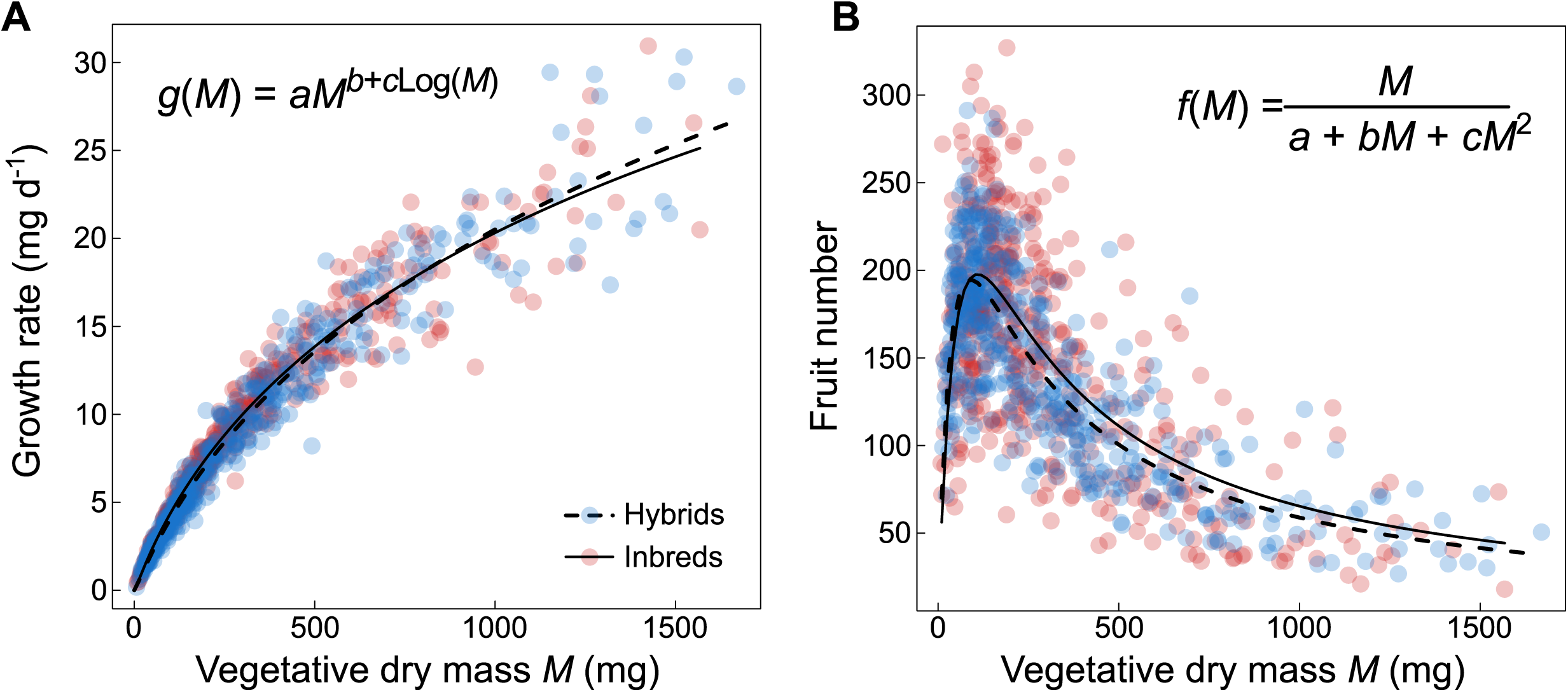
Allometric relationships of growth rate and fruit number. (**A**) Allometric relationship between growth rate and vegetative plant dry mass, *M*, fitted with a power-law function with mass-corrected exponent (*g*(*M*)). Equation fitted separately on 451 inbreds (red dots, solid line) and 447 hybrids (blue dots, dashed line). (**B**) Allometric relationship between fruit number and *M*, fitted with an inverse polynomial function (*f*(*M*)). Equation fitted separately on 441 inbreds (red dots, solid line) and 449 hybrids (blue dots, dashed line).

### Relationships between heterosis and genetic, phenotypic and geographic distances

We first compared what fraction of heterosis could be explained by genetic and phenotypic distances between inbred parents. For all hybrids, we quantified heterosis as the observed phenotypic deviation relative to mid-parent value (MPH), and best parental value (BPH). Pairwise genetic distances were calculated either with all SNPs in the genome, or with SNPs in the 1% top-genes associated with the corresponding trait [27]. For the phenotypic distance, we used the absolute difference in vegetative dry mass between parents.

We found that genetic distance between parents was positively correlated with heterosis of growth rate (both MPH and BPH *P* < 0.001; Fig. 5A). However, genetic distance only accounted for less than 10% of heterosis (7% and 6% for MPH and BPH, respectively). Phenotypic distance was also poorly correlated with heterosis of growth rate, and only with MPH (*r*^2^ = 0.03, Fig. 5B). By contrast, genetic distance did not correlate with heterosis of fruit number (both P > 0.01, Fig. 5C), while phenotypic distance was negatively correlated, albeit poorly, with BPH (*r*^2^ = 0.05, Fig. 5D). Using only the 1% top-genes associated with growth rate and fruit number, which are more likely to have a causal role in the measured traits, did not improve correlations (*r*^2^ < 0.10). Even when combined in a multivariate model, genetic and phenotypic distances together explained less than 10% of heterosis of growth rate and fruit number (Table S3). For comparison, geographic distance between parents explained less than 2% of heterosis of fruit number, and less than 0.2% of heterosis of growth rate. In summary, none of the parental distances – be they genetic, phenotypic or geographic – had much power to explain and predict a significant portion of heterosis.

**Figure 5.**
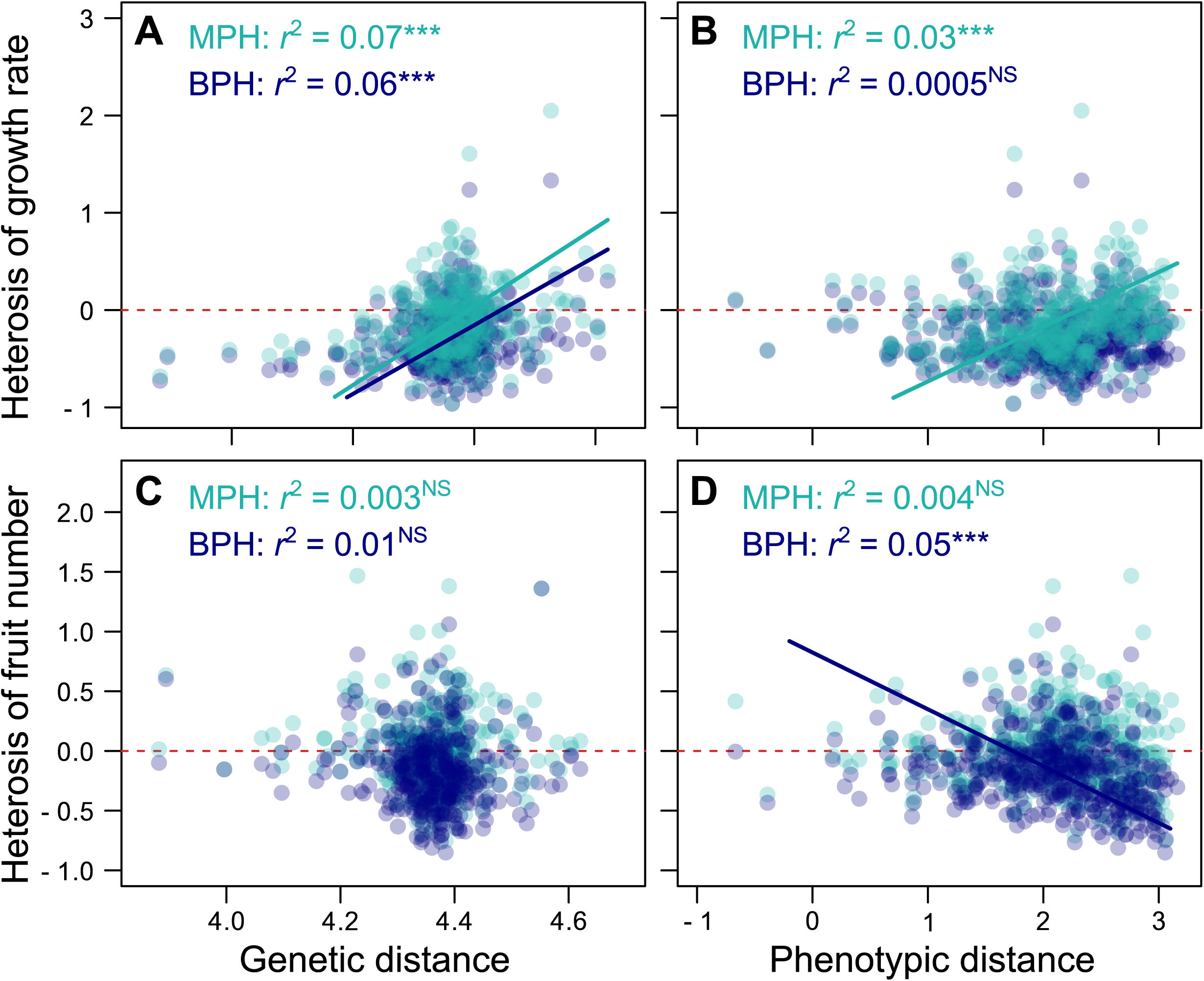
Correlation of heterosis with genetic and phenotypic distances. (**A**) Correlation between pairwise genetic distances between parental accessions and observed heterosis of growth rate in hybrids (*n* = 368). (**B**) Correlation between pairwise phenotypic distances between parental accessions (absolute difference in *M*) and observed heterosis of growth rate in hybrids (*n* = 447). (**C**) Pairwise genetic distances versus heterosis of fruit number (*n* = 368). (**D**) Pairwise phenotypic distances versus heterosis of fruit number (*n* = 449). MPH: Mid-Parent Heterosis (light blue), BPH: Best-parent heterosis (dark blue). *r*^2^ are Pearson’s coefficients of correlation (NS: Non-significant, ***: *P* < 0.001). Regressions lines are SMA, drawn when significant (solid lines). Red dashed lines represent zero axis.

### Explaining variation in heterosis by phenotypic non-linearity

In a second approach, we used the fitted equations (Table 1) to take into account the non-linearity of allometric relationships and to predict heterosis. Our first goal was to predict growth rate and fruit number with (i) the allometric relationship fitted on parents, and (ii) the measurement of vegetative dry mass in hybrids (Fig. 6). Predicted growth rate in hybrids strongly correlated with observed growth rate (*r*^2^ = 0.95, Fig. S1A). By contrast, predicted fruit number was poorly correlated with observed trait values (*r*^2^ = 0.08, Fig. S1B). This is consistent with the larger dispersion of trait values around the fitted curve for fruit number (Fig. 4B) compared to growth rate (Fig. 4A). We then compared trait deviation predicted in hybrids by non-linearity of traits – relative to the predicted mid-parent value (PNL_MP_) and to the predicted best-parent value (PNL_BP_) – with the observed MPH and BPH (Fig. 6). Observed heterosis and predicted non-linearity of growth rate were strongly correlated (*r*^2^ = 0.75 and 0.66 for PNL_MP_ vs MPH and PNL_BP_ vs BPH, respectively, Fig. 6B). Observed heterosis and predicted phenotypic non-linearity of fruit number were also positively correlated, although weaker than for growth rate (*r*^2^ = 0.14 and 0.10 for PNL_MP_ vs MPH and PNL_BP_ vs BPH, respectively, Fig. 6D). This suggests that allometric relationships allow the prediction of heterosis amplitude, and that prediction accuracy depends on the strength of the underlying non-linear covariation between traits.

**Figure 6.**
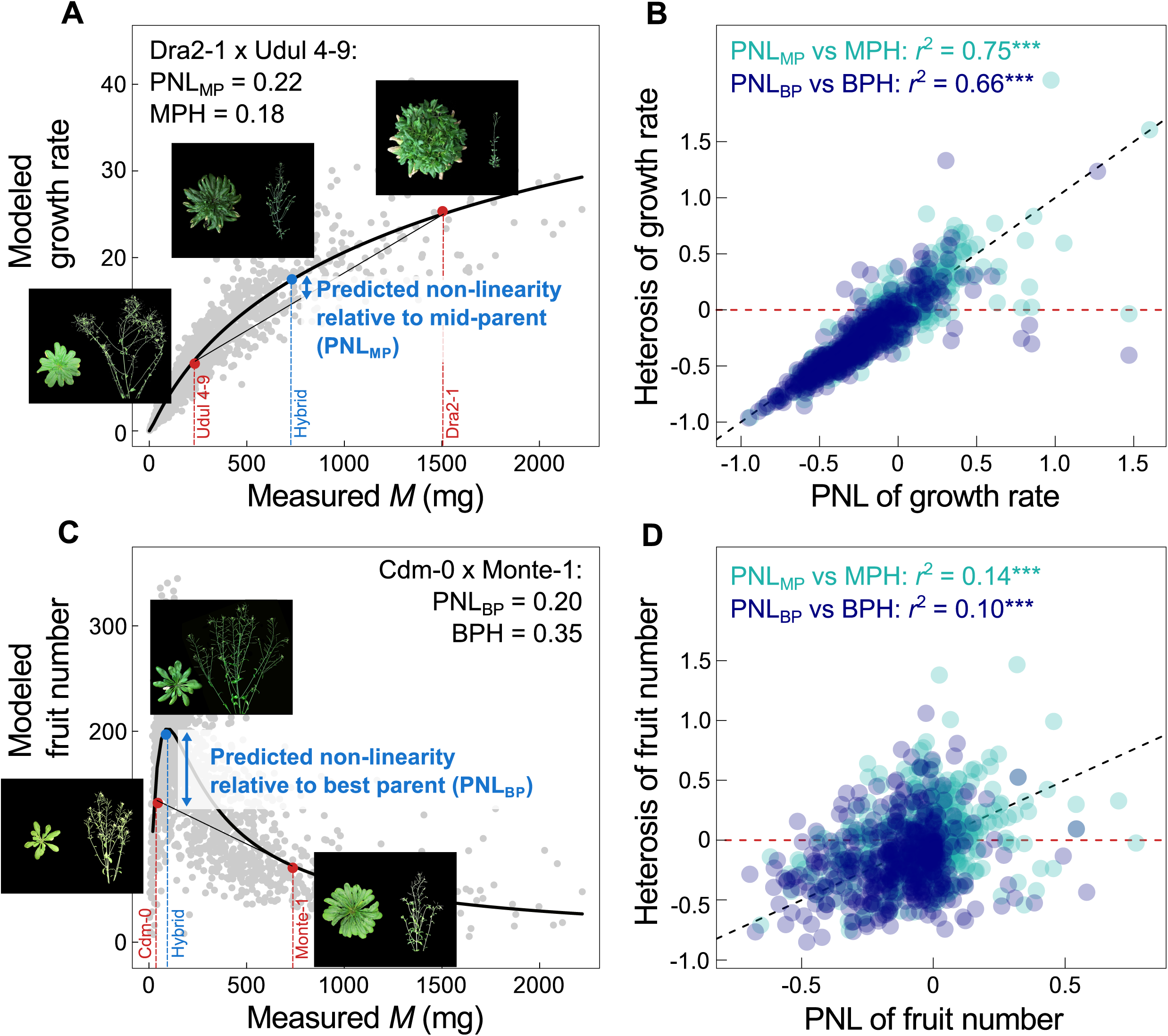
Prediction of heterosis with phenotypic non-linearity. (**A**) Allometric relationship of growth rate (*g*(*M*), black solid line) fitted in inbreds, with an example of two parental accessions (Udul 4-9 and Dra2-1, in red), and their hybrid (in blue). Phenotypic non-linearity (PNL, blue arrows) was calculated relative to mid-parent value (MP_pred_): PNL_MP_ = (*g*(*M*_1×2_) - MP_pred_) / MP_pred_, and relative to best-parent value (BP_pred_): PNL_BP_ = (*g*(*M*_1×2_) - BP_pred_) / BP_pred_. (**B**) Correlation between PNL and heterosis in all hybrids (*n* = 447), both relative to mid- and best-parent value (MPH in light blue and BPH in dark blue, respectively). (**C**) Allometric relationship of fruit number (*f*(*M*), black solid line) fitted in inbreds, with an example of two parental accessions (Cdm-0 and Monte-1, in red), and their hybrid (in blue). PNL was calculated relative to mid-parent value (PNL_MP_) and relative to best-parent value (PNL_BP_). (**D**) Correlation between PNL and heterosis in all hybrids (*n* = 447), both relative to MPH (light blue) and BPH (dark blue). Red dashed lines represent zero axes. Black dashed line represents 1:1 line. Grey dots are all hybrid individuals. Pictures of plants were taken at harvesting, at the end of reproduction. *r*^2^ are Pearson’s coefficients of correlation (NS: Non-significant, ***: *P* < 0.001).

## Discussion

Already in 1934, Wright wrote in his seminal paper that “dominance has to do with the physiology of the organism and has nothing to do with the mechanism of transmission” [38]. Eighty years later, the emergence of heterosis is still considered as an enigma and its physiological bases remain debated. Despite the many dominant, overdominant and epistatic QTL identified in a plethora of species [19,20,36], none of the genetic models has been formally validated in more than a few cases [*e.g.*, 9]. Here, we approached the question of heterosis from a physiological angle based on a geometric analysis of trait-trait relationships. Combining the ideas put forth by Wright [38] with principles from allometric theory [44,51], we demonstrate that a significant part of heterosis can be explained by the non-linear relationships that link phenotypic traits with each other.

The accuracy of heterosis prediction with phenotypic non-linearity depends on how strongly traits correlate with each other. Metabolic scaling theory [44,46,66] postulates that body size constrains trait variation in the form of universal mathematical laws. According to the postulates of this theory, biomass determines the number of metabolically active units (cells, mitochondria), which in turn determine physiological fluxes, biomass partitioning and growth rate at the organismal level [51,67]. In agreement, we have found that hybrids exhibit the same allometric relationships as natural inbred accessions. This reinforces the idea of strong and predictable constraints on phenotypic variation, specifically on variation of growth rate. Our results further indicate that the shape of phenotypic non-linearity is important for the emergence of heterosis [40]. For instance, a severe concavity is expected to be favourable to the emergence of best-parent heterosis, as we have observed for fruit number. By contrast, phenotypic convexity with a trait at lower integration level, for instance a trait related to organism development and organization plan, could potentially lead to negative heterosis. This could notably explain the high fraction of negative heterosis observed here for plant biomass, which is consistent with previous findings in *A. thaliana* [13,54], although it contrasts with some others [7,56]. It is also consistent with results in rice [17,68], an inbreeding crop species used for the development of heterotic hybrids. In addition to phenotypic convexity, another explanation to the emergence of negative heterosis is that we used crosses from strongly divergent accessions, which might exhibit incompatibilities between defence-related genes [69-72]. Indeed, autoimmunity has been repeatedly observed when crossing distant accessions of *A. thaliana* [64]. This is also consistent with recent findings, where positive heterosis for biomass seems to be linked to a general repression of the immune system in *A. thaliana* hybrids [56].

The genetic bases of fitness-related traits such as growth rate and fruit production are complex by nature, because these traits result from the effects of numerous genes acting on different components of performance [26,73]. Inexpensive high-density genotype information coupled with very detailed phenotyping provides a series of promising avenues for the genomic prediction of heterosis [74]. In this context, the dominance hypothesis implicates, within certain limits, a positive relationship between parental genetic distance and heterosis [28,29]. Results from a range of species, however, do not conform with these expectations [7,13,34,75,76]. One of the reasons could be the often small number of crosses and the relatively small range of genetic distances analysed, with the latter holding true especially in cultivated species [7]. Our results with 450 hybrids representing crosses between diverse *A. thaliana* populations pointed to a positive but weak correlation between heterosis and parental genetic distance for growth rate, but no such correlation for fruit number. This suggests that a genetic approach alone may not be sufficient to accurately predict heterosis. Importantly, genetic distances calculated from the genes enriched for the ones likely to have major effect on growth rate [27] did not improve our ability to predict heterosis.

Wright’s model of physiological dominance [38] was based on the non-linear relationship that connects two traits at different levels of integration, enzyme concentration and metabolic fluxes. In evolutionary biology, a trait is said to be “integrated” when it is closely associated with the fitness of the organism, which in plants typically includes growth rate and fruit number [77]. This leads to a pyramidal view of phenotypic integration [77,78], with fitness-related traits at the top being under the control of several, less integrated traits (*e.g.*, biomass, phenology, morphology), which themselves are under the control of many other traits (*e.g.*, metabolic and physiological fluxes, cellular activity), and so on, until traits supposed to be additive such as enzyme concentrations. Under the hypothesis of additivity of enzyme concentration, Wright’s model has been shown to accurately predict heterosis of metabolic output in the chain of glycolysis [40]. In our study, we have shown that even when vegetative dry mass exhibited important deviation from additivity, it could still be used to predict heterosis of growth rate and fruit number based on parental allometric relationships. This finding demonstrates the performance of the phenotypic approach relative to the purely genetic approach to explain and predict heterosis. For instance, our method explained up to 75% of heterosis amplitude for growth rate (Fig. 6B) while genetic distance at best explained only 7% (Fig. 5A). Although our approach is not incompatible with the genetic models of molecular dominance, it outperforms these in its ability to predict the direction and amplitude of heterosis in the natural inbred species *A. thaliana*.

Evolutionary theory suggests that the intensity of inbreeding depression, and hence the potential for heterosis, should increase with the level of phenotypic integration [79]. Consistent with this, survival and fertility in animals are more sensitive to inbreeding depression than traits at lower levels of integration such as size, biomass and gross morphology [80]. Our findings are consistent with the idea that fitness-related traits exhibit more heterosis because they result from the multiplicative effects of non-linear relationships at different levels of phenotypic integration [48,73,81]. These non-linear relationships can have different curvatures and directions, which might either amplify or cancel each other.

From a theoretical point of view, the predominance of outcrossing and the maintenance of recessive alleles among organisms have been suggested to be directly linked to non-additive inheritance and superior performance of hybrids [82]. Wright’s initial model of physiological dominance was proposed as a response to Fisher’s idea that ‘gene modifiers of dominance’ must exist and be selected to maximize the fitness of the heterozygote [83]. Wright [38], and later Kacser and Burns [39], claimed that modifiers are not necessary, because non-linearity is an intrinsic characteristic of metabolic networks. The same argument holds true for complex traits such as growth rate and fruit number. The assumption of modifiers of dominance is based on the unrealistic expectation of an intermediate phenotype in the heterozygote, while phenotypic relationships are essentially non-linear [40,48,84]. A next step in getting to the physiological root of heterosis will be the integration of other, more low-level traits, especially gene expression and metabolite levels. We expect that this will improve the characterization of non-linear relationships in multidimensional phenotypic space, and ultimately shed lights on the physiological mechanisms at the origin of non-additive inheritance.

## Conclusions

The development of a predictive approach for heterosis represents a long-term goal of modern biology, especially in the applied framework of varietal selection in crops [85]. Our study highlights the power of a geometric approach of trait-trait relationships for explaining heterosis in two major components of plant productivity and yield. This in turn opens avenues for targeting optimal crosses based on allometric relationships in parental lines. That trait variation is similarly constrained in accessions and hybrids suggests that hybrids outperforming all accessions in a specific trait are the exception, although they generally outperform their specific parents. It is now time to test the phenotypic approach to heterosis in cultivated species, for which the study of allometric relationships is a nascent research front [86–88].

## Material and Methods

### Plant material

451 accessions of *Arabidopsis thaliana* were phenotyped in 2014 at the Max Planck Institute for Developmental Biology in Tübingen (MPI-Tübingen), Germany (*n* = 2, Exp1) [27,89]. 415 accessions were used as parental lines of the 450 hybrids phenotyped in a second experiment in 2015 at MPI-Tübingen (*n* = 4, Exp2). 342 accessions were used as female parent and 318 as male parent. 407 accessions among the 451 total, and 369 accessions among the 415 parental, had been genome sequenced as part of the 1001 Genomes project (http://1001genomes.org/)[59]. Among the 415 parental accessions, 134 (32%) were used in a single cross, 166 (40%) were used in two crosses, 63 (15%) in three crosses, and 52 (13%) in at least four crosses. To overcome potential maternal effects, the same mother plants grown in the greenhouse in 2013 provided the seeds for both the inbreds (by self-fertilization) and the hybrids (by manual cross) (Fig. S2A).

### Growth conditions

We designed a hydroponic system where plants were cultivated on inorganic solid media (rockwool) and all nutrients were provided through the watering solution. 4.6 cm (diameter) x 5 cm (depth) circular pots (Pöppelmann, Lohne, Germany) were filled with 3.6 cm x 3.6 cm x 4 cm depth rockwool cubes (Grodan cubes, Rockwool International, Denmark). Pots were covered with a black foam disk with a 5-10 mm central circular opening. Seeds were sown in individual pots, randomly distributed in trays of 30 pots each (Fig. S2B).

Before sowing, all seeds were surface-sterilized with 100% ethanol and frozen overnight at −80 °C to kill any insect eggs. The rockwool cubes were placed in 75% strength nutrient solution as described in ref. [90] in order to achieve full humidification and fertilization. After sowing on the surface of the rockwool cubes, trays with 30 pots each were incubated for two days in the dark at 4°C for stratification. Trays were transferred for 6 days to 23°C (8 h day length) for germination. After 6 days, when most seedlings had two cotyledons, trays were transferred to 4°C (8 h light) for 41 days of vernalization, in order to reduce the range of flowering times among accessions. During germination and vernalization, all trays were watered once a week with a 75% strength nutrient solution. After vernalization, when true leaves had emerged on most individuals, plants were thinned to one individual per pot, and trays were moved to the Raspberry Pi Automated Plant Analysis (RAPA) facility [89] (Fig. S2B), set to 16°C, air humidity at 65%, and 12 h day length, with a PPFD of 125 to 175 µmol m^-2^ s^-1^ provided by a 1:1 mixture of Cool White and Gro-Lux Wide Spectrum fluorescent lights (Luxline plus F36W/840, Sylvania, Germany). Trays were randomly positioned in the room, and watered every 1 to 3 days with 100% strength nutrient solution.

### Trait measurement

Plants were grown and phenotyped using rigorously the same protocol, following methodologies previously published for inbred accessions [27,89]. Plants were harvested at the end of the life cycle when the first fruits were senescing. Rosettes were separated from roots and reproductive parts, dried at 65° C for three days and weighed. Plant lifespan (d) was measured as the duration between the appearance of the two first leaves after vernalization and the end of the life cycle [89]. Growth rate (mg d^-1^) was calculated as the ratio of final rosette dry mass over plant lifespan. Inflorescences were photographed with a high-resolution, 16.6 megapixel SLR camera (Canon EOS-1, Canon Inc., Japan), and analysed with ImageJ [91] to estimate the number of fruits through 2D image skeletonization, following published protocols [27,89].

The RAPA system was used for daily imaging using 192 micro-cameras (OmniVision OV5647), which simultaneously acquired 6 daily top-view 5-megapixel images for each tray during the first 25 days after vernalization. We used a published method to estimate plant dry mass during ontogeny from top-view rosette pictures [27,89]. From fitted sigmoid growth curves on all individuals, we calculated inflection point (d) at which daily growth was maximal and started to decrease. We used rosette dry mass (mg) at the inflection point as measurement of vegetative dry mass *M* (mg). In total, trait values for plant lifespan, growth rate and vegetative dry mass were available on 451 inbreds and 447 hybrids, and fruit number on 441 inbreds and 449 hybrids (Table S1).

To correct for potential biases between the two experiments performed, 16 accessions phenotyped in Exp1 were also included in Exp2. Among all traits measured, only plant lifespan exhibited a significant difference between the two experiments (*P* = 0.03). We thus corrected trait values in Exp2 with the following equation: Lifespan_corrected_ = −37 + 1.8*Lifespan_observed_.

### Measurement of heterosis

First, hybrid phenotypic classes were categorized by comparing trait distribution in hybrids to mean and best (or worst) parental values. We used 95% confidence intervals (CI) among the four hybrid replicates of each F1 for categorization, as follows: (i) below worst-parent if CI of the trait measured in the hybrid was strictly inferior to the minimum trait value among the two parental genotypes; (ii) below mean-parent if the hybrid CI was strictly inferior to mean parental value but overlapped with minimum parent; (iii) above mean-parent if the hybrid CI was strictly superior to mean parental value (but not best parent); (iv) above best-parent if the hybrid CI was strictly superior to maximum parental value.

Secondly, two metrics commonly used in the literature to quantify heterosis [13] were calculated in the present study for all traits, Y, *i.e.* growth rate and fruit number:

1. Mid-Parent Heterosis (MPH), the deviation of observed hybrid value scaled to observed mean parental value:

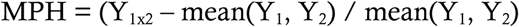
2. Best-Parent Heterosis (BPH), the deviation of observed hybrid value scaled to observed best parental value:

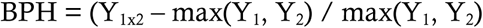

### Allometric relationships and measurement of phenotypic non-linearity

The allometric equations were fitted by the non-linear least-squares method using the *nls* function in R [92]. Based on allometric theory [27,43,44,52], we chose a power-law equation for growth rate with a mass-corrected allometric exponent (*g*(*M*) in Table 1). The corrected exponent corresponds to the derivative of the quadratic function obtained after logarithmic transformation of the allometric relationship [27,43,52]. In our data, biomass-correction of the exponent improved the fitting of the allometric relationship of growth rate (ΔAIC = 83). For the allometric relationship of fruits number, we chose an inverse polynomial equation (*f*(*M*) in Table 1). 95% CIs of the fitted coefficients were estimated with the *confint* function in R.

We used the allometric equations fitted on the accessions to predict the phenotype of the hybrids from vegetative dry mass *M*. We first estimated growth rate of both parents and hybrids (*g*(*M*_1_), *g*(*M*_2_), *g*(*M*_1×2_) with *M*_1_ and *M*_2_ corresponding to vegetative dry mass of worst and best parent, respectively, and *M*_1×2_ to hybrid dry mass). We then predicted mean parental value of growth rate as MP_pred_ = (*g*(*M*_1_) + *g*(*M*_2_))/2, and best parental value as BP_pred_ = max(*g*(*M*_1_), *g*(*M*_2_)). Finally, we predicted the phenotypic non-linearity as:

1. the deviation of predicted hybrid value scaled to predicted mean parental value (Fig. 6A):

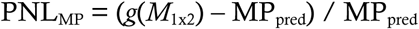
2. the deviation of predicted hybrid value scaled to predicted best parental value (Fig. 6C):

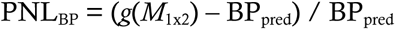

Both PNL_MP_ and PNL_BP_ were calculated for growth rate, and we performed similarly for estimating the non-linearity of fruit number (Fig. 6C).

### Genetic, geographic and phenotypic distances

407 accessions out of the 451 phenotyped here have been genome sequenced (http://1001genomes.org/) [59]. Using vcftools [93], the 12,883,854 single nucleotide polymorphisms (SNPs) were first filtered to retain those where minor allele frequency was above 5%, with a genotyping rate above 85% across all accessions. This resulted in 391,016 SNPs. We used PLINK v1.9 [94] to estimate pairwise genetic distances as the number of alleles that differed between pairs of accessions (*--distance* function), after log_10_-transformation. We also measured genetic distances using the 1% SNPs with the strongest (positive and negative) effects on growth rate or fruit number from a previously published study [27]. SNPs effects on each trait were estimated using a polygenic GWA model implemented in GEMMA (‘Bayesian Sparse Linear Mixed Model’, BSLMM) [95].

Pairwise geographic distances between accessions were estimated from their longitude-latitude coordinates [59] and the *distm* function of the geosphere package in R. For the calculation of pairwise phenotypic distances, we first used Euclidean distance among all traits measured in inbreds (vegetative dry mass, lifespan, growth rate, fruit number; used for Fig. 1). We also used absolute difference in vegetative dry mass between parents for comparing the contribution of genetic and phenotypic distances to heterosis in Fig. 5.

### Statistical analyses

Pearson’s method was used for correlation analyses (*cor.test* function in R). Linear regressions were drawn with Standard Major Axis (SMA, ‘smatr’ package in R). The effect of the experiment on trait values was tested on the 16 accessions common to Exp1 and Exp2, using two-way ANOVA with genotype and experiment as interacting factors. 95% Confidence Interval (CI) was calculated as 1.96*Standard Deviation. To calculate the proportion of phenotypic variance associated with genotypic (G) effects (a measure of broad-sense heritability, *H*^2^ = var(G) / (var(G) + residuals)), we fitted a mixed model (*lmer* function in R) as Y = Genotype + residuals, where Y is trait value and Genotype is used as random factor. Statistical analyses were conducted in R v3.2.3 [92].

### Availability of data and material

The datasets supporting the conclusions of this article are included within the article and its additional files. Codes are available on Github (https://github.com/fvasseur/AnalysisHeterosis), and phenotypic data are available on the Dryad repository (https://datadryad.org/review?doi=doi:10.5061/dryad.4978rp4) [96]. Correspondence and requests for materials should be addressed to.

## Author Contributions

FV, CV and DW formulated hypotheses and designed the study. FV performed the experiments and extracted the data. FV, LF and DV performed statistical analyses. All authors interpreted the results and wrote the paper.

## Acknowledgements

We thank Christine Dillmann, Alain Charcosset, Olivier Loudet and Thomas Lenormand for helpful comments about statistical modelling and data interpretation.

## Additional Files

**Additional Table S1. Traits measured in inbred accessions and hybrids of *A. thaliana***

**Additional Table S2. Summary statistics of traits measured.**

**Additional Table S3. Correlation of heterosis with genetic and phenotypic distances.**

**Additional Figure S1. Correlation between predicted and observed hybrid phenotype.** (**A**) Hybrid growth rate (mg d^-1^) predicted by phenotypic non-linearity (brown dots) and genetic additivity (mean growth rate between parents, yellow dots), and compared to observed hybrid value (*n* = 447). (**B**) Hybrid fruit number predicted by phenotypic non-linearity (brown dots) and genetic additivity (mean fruit number between parents, yellow dots), and compared to observed hybrid value (*n* = 449). *r*^2^ are Pearson’s coefficients of correlation (NS: Non-significant, ***: *P* < 0.001). Dashed line represents 1:1 line.

**Additional Figure S2. Crossing and phenotyping conditions.** (**A**) Seed production experiment performed in 2013 at MPI-Tübingen (Germany). After manual crossing or self-fertilization, mother flowers were isolated in a small paper bags until fruit ripening. Seeds for accessions and hybrids used in this study came from the same mother plants. (**B**) RAPA growth chamber at MPI-Tübingen with trays of accessions phenotyped during Exp1 in 2014.

